# Stopover Population Estimate and Migration Ecology of Red Knots *C. c. rufa* at Delaware Bay, USA, 2025

**DOI:** 10.64898/2026.02.25.708011

**Authors:** J. E. Lyons

## Abstract

Red Knots (*Calidris canutus rufa*) rely on Atlantic horseshoe crab (*Limulus polyphemus*) eggs in the Delaware Bay to refuel during northward migration. Intensive harvest of horseshoe crabs in the 1990s contributed to declines in Red Knot numbers. In 2013, the Atlantic States Marine Fisheries Commission adopted an Adaptive Resource Management (ARM) framework to balance sustainable horseshoe crab harvest with ecosystem integrity and Red Knot recovery, requiring annual stopover population estimates. We estimated the 2025 passage population of Red Knots at Delaware Bay using a Bayesian analysis of a Jolly–Seber mark–resight model which accounts for population turnover and imperfect detection. We also evaluated change in migration timing between 2011 and 2025 with model-derived estimates of arrival at the Delaware Bay each year. The 2025 passage population was 54,043 individuals (95% credible interval: 47,926–61,928), an increase of approximately 17% over 2024 and only the second year since 2011 to exceed 50,000 individuals. Despite the increase, overlapping credible intervals across years indicate a stable stopover population. Migration timing has remained consistent, with 50% of the population typically arriving by 18 May and no evidence of advancement since 2011. These findings provide meaningful input for the ARM framework, supporting sustainable harvest of horseshoe crabs while maintaining adequate foraging opportunities for Red Knots and other shorebirds.

*Parts of the Introduction, Methods, and Appendices were originally published in Lyons (2024) and are summarized herein*.

## 1 Introduction

The northward migration of Red Knots (*Calidris canutus rufa*) in the mid-Atlantic region coincides with the onset of spawning of horseshoe crabs (*Limulus polyphemus*). Red Knots stop at the Delaware Bay to feed on horseshoe crab eggs, which are an important food resource for Red Knots and other shorebirds because they have a high energy content and are easily digestible (Karpanty et al., 2006, Haramis et al., 2007).

Horseshoe crabs have been harvested since at least 1990 for use as bait in American eel (*Anguilla rostrata*) and whelk (*Busycotypus canaliculatus* and *Busycon carica*) fisheries (Kreamer and Michels, 2009). The number of horseshoe crabs harvested increased throughout the 1990s, peaked in the late 1990s, and then declined in the early 2000s. In the 1990s and early 2000s the estimated number of Red Knots counted during aerial surveys of Delaware Bay declined from ∼50,000 to ∼13,000 (Niles et al., 2008). Avian conservation biologists hypothesized that unregulated harvest of horseshoe crabs in Delaware Bay during the 1990s reduced food availability for migratory shorebirds. This shortage limited their ability to refuel during stopover, thereby potentially compromising migration performance, breeding success, and annual survival (Baker et al., 2004, McGowan, Hines, et al., 2011).

The Atlantic States Marine Fisheries Commission (ASMFC) has managed the horseshoe crab harvest in the Delaware Bay region since 1998 and in 2012 adopted an Adaptive Resource Management (ARM) framework that was created to explicitly incorporate shorebird objectives in horseshoe crab harvest management (McGowan, Smith, et al., 2015). The ARM framework was designed to constrain the harvest so that the number of spawning horseshoe crabs would not limit the number of Red Knots stopping at Delaware Bay during migration. To achieve multiple objectives simultaneously, the ARM framework requires an estimate each year of both the horseshoe crab population and the Red Knot stopover population (McGowan, Lyons, and Smith, 2015).

The timing of arrival at key stopover sites and stopover duration drive stopover dynamics. Variability in arrival timing can influence how effectively birds access food resources and may affect survival and subsequent breeding success (Baker et al. 2004, Atkinson et al. 2007). Moreover, shifts in migration timing can signal broader ecological changes, such as climate-driven alterations to food availability or habitat conditions (Visser et al. 2012). Documenting and analyzing arrival patterns over time therefore provides valuable insight into population dynamics and helps guide management actions aimed at maintaining the ecological integrity of vital stopover habitats.

The goals of this study were to 1) estimate the stopover population size in 2025, as we have each year since 2011, using mark-resight data on individually marked birds and a Jolly-Seber model for open populations (Jolly, 1965, Seber, 1965) to support the ARM framework, and 2) assess potential shifts in migration timing since 2011 using model-based estimates of arrival at Delaware Bay.

## 2 Methods

### 2.1 Field methods

Since 2003, Red Knots have been individually marked at Delaware Bay and other regions across the Western Hemisphere (e.g., parts of Argentina, Brazil, Canada, Chile) with field-readable leg flags engraved with unique three-character alphanumeric codes (Clark et al., 2005). Mark–resight data consisting of individual resightings of flagged birds and counts of marked and unmarked birds were collected during northward migration in 2025 on the Delaware and New Jersey shores of Delaware Bay following established protocols (Lyons, 2016). These methods have been consistently applied at Delaware Bay since 2011.

Surveys were conducted on 20 beaches (Appendix A) during May and early June 2025 (Table 1). Each beach was surveyed at least once in every three-day sampling period (Lyons, 2016). During surveys, agency staff and volunteers searched for flagged birds, recording the engraved alphanumeric codes. During resighting surveys, observers also periodically employed scan sampling techniques to estimate the marked–unmarked ratio. In each scan, a random portion of a Red Knot flock was examined, and observers recorded (1) the number of individually flagged birds and (2) the total number of birds checked for flags (Lyons, 2016). Survey data are available from Delaware Division of Fish and Wildlife and New Jersey Department of Environmental Protection.

**Table 1:** Dates for mark-resight survey periods (3-day sampling occasions) for Red Knot (*C. c. rufa*) population analysis at Delaware Bay in 2025. The same sampling periods have been used at Delaware Bay since 2011.

| Survey period | Dates | Survey period | Dates |
| --- | --- | --- | --- |
| 1 | 8–10 May | 6 | 23–25 May |
| 2 | 11–13 May | 7 | 26–28 May |
| 3 | 14–16 May | 8 | 29–31 May |
| 4 | 17–19 May | 9 | 1–3 June |
| 5 | 20–22 May | 10 | 4–6 June |

### 2.2 Statistical methods

For analysis with the Jolly-Seber (JS) model, the migration season at Delaware Bay was divided into 10 three-day sampling periods (Table 1). Three days was chosen as the time step for the mark-resight model given the time required to complete a circuit of all 20 beaches. Multiple observations of a flagged individual within any of these sampling periods were combined into one observation per period and formatted as encounter histories for mark-recapture modeling (Williams, Nichols, and Conroy, 2002). A summary of the 2025 mark–resight data is provided in Appendix B.

Stopover population size was estimated using methods outlined in Lyons et al. (2016). Specifically, we used the superpopulation framework of Crosbie and Manly (1985) and Schwarz and Arnason (1996) for the JS model (Jolly, 1965; Seber, 1965), which is an open-population model accounting for turnover of individuals during migration and imperfect detection during surveys. The superpopulation is defined as all birds present in the study area on at least one sampling occasion (Nichols and Kaiser, 1999).

When applied at migration stopover sites, the JS model includes time-specific parameters for (1) probability of arrival to the study area, (2) probability of departure, (3) probability of detection (resighting) during a sampling occasion, and (4) the total passage population size, or superpopulation. Resighting probability was modeled as a function of flag color because some colors are harder to read than others (Appendix C); specifically, we fit a separate detection probability for dark green flags because these can have lower detection probability than light green flags (Tucker, McGowan, Nuse, et al., 2023). Departure probability is the probability that a bird present at Delaware Bay during sampling period *i* departs before sampling period *i* + 1. Arrival, departure, and resighting parameters are biologically meaningful at Delaware Bay because they capture the dynamic nature of migration, where birds arrive and depart continuously, remain for varying durations, and are resighted with imperfect detection, allowing estimation of the total number of Red Knots using the stopover (superpopulation).

We used a Bayesian analysis of a state-space formulation of the JS model (Royle and Dorazio, 2008; Kéry and Schaub, 2012; Lyons et al., 2016) in which abundance estimates were derived in two steps. First, the number of birds with leg flags was estimated using encounter histories. Second, this abundance estimate was adjusted for unmarked birds using the proportion of flagged individuals (Lyons et al., 2016). The flagged proportion was estimated from the scan sample data using a binomial model, implemented as a generalized linear mixed model with a random effect for sampling period to account for temporal variation in arrival of flagged birds and additions of newly-flagged individuals during the migration season. Additional methodological details are provided in Lyons et al. (2016) and Appendix C.

The JS model has several key assumptions, including homogeneous probabilities of arrival, stopover departure, and resighting among individuals; that marks are not lost or misrecorded; that sampling is effectively instantaneous relative to the interval between occasions; that emigration from the study area is permanent; that individual fates are independent of one another; and that marked-to-unmarked ratio data are representative of the population (i.e., individuals in the scan samples are randomly selected from the population). To help meet the assumption that individual identities of flagged birds were recorded without error, resighting data were validated against physical capture and flag-deployment records in the http://www.bandedbirds.org/ repository. Resightings without a corresponding record of prior physical capture and flag deployment were excluded from analyses as false-positive (misread) errors. An exception was made for resightings of orange flags, which were retained without validation because capture records from Argentina and Uruguay—where orange flags are used under the Pan American Shorebird Program (Howes, Béraud, and Drolet-Gratton, 2016)—are generally not available in the repository. Thus, validation against capture data was applied when such data were available but could not be applied uniformly across all countries. A second type of false positive error is also possible. Some of the resightings in our dataset may be flags that were deployed prior to 2025 but were not in fact present at Delaware Bay in 2025. It is not possible to identify this second type of false positive with banding data validation or other quality assurance/quality control methods (Tucker, McGowan, Robinson, et al., 2019). See Lyons (2016) for discussion of all model assumptions.

Stopover duration is an important reflection of turnover and key aspect of stopover dynamics. We estimated stopover duration in 2025 using the approach described by Lyons et al. (2016) and Lok et al. (2019), which takes advantage of the state-space formulation of the Jolly–Seber model. In the state-space formulation, a latent parameter indicates the number of time steps that each individual remains in the study area, i.e., stopover duration. Lyons et al. (2016) originally calculated duration based only on individuals that were resighted at least once. Lok et al. (2019) improved this approach by extending the calculation to all individuals estimated to be present for at least one sampling period, including those never observed, thereby reducing bias toward longer stopovers that can result from excluding short-stay individuals.

To assess potential temporal shifts in arrival timing since 2011, we calculated the date by which 50% of the population had arrived each year from 2011 to 2025. For each of 10,000 posterior draws from the arrival probability parameters, we computed the cumulative sum of arrivals across sampling occasions (Table 1) within each year. The 50% arrival date was defined as the first occasion where the cumulative proportion reached or exceeded 0.5. We summarized the posterior distribution of 50% arrival dates using median values and 95% credible intervals for each year. Temporal trends were evaluated using linear regression of annual median arrival occasion against year, with statistical significance assessed at *α* = 0.05. We also calculated the proportion of posterior draws reaching 50% arrival before occasion 4 (18 May) as an indicator of early arrival years. This approach accounts for both within-year uncertainty in arrival estimates and between-year variation in the timing of the 50% threshold.

## 3 Results

### 3.1 Mark-resight surveys

Field observers in 2025 collected 2,035 flag resightings, including multiple observations per flag in different sampling periods. One-hundred and nineteen (119) resightings were not valid (i.e., no corresponding banding data), for an overall misread error of 5.8%. The valid resightings represented a total of 1,276 individual birds that were recorded at least once between 7 May and 6 June 2025. On average, each flagged individual was resighted 1.5 times and the maximum number of times resighted was eight. These individuals were originally captured and flagged in five to seven different countries (Fig. 1). The number of flagged individuals, i.e., encounter histories for analysis, in 2025 was slightly lower than in 2024 (1,395) and the second lowest number of flagged individuals detected since 2011; only 2023 had fewer flags (1,091; Fig. 2).

**Figure 1.**
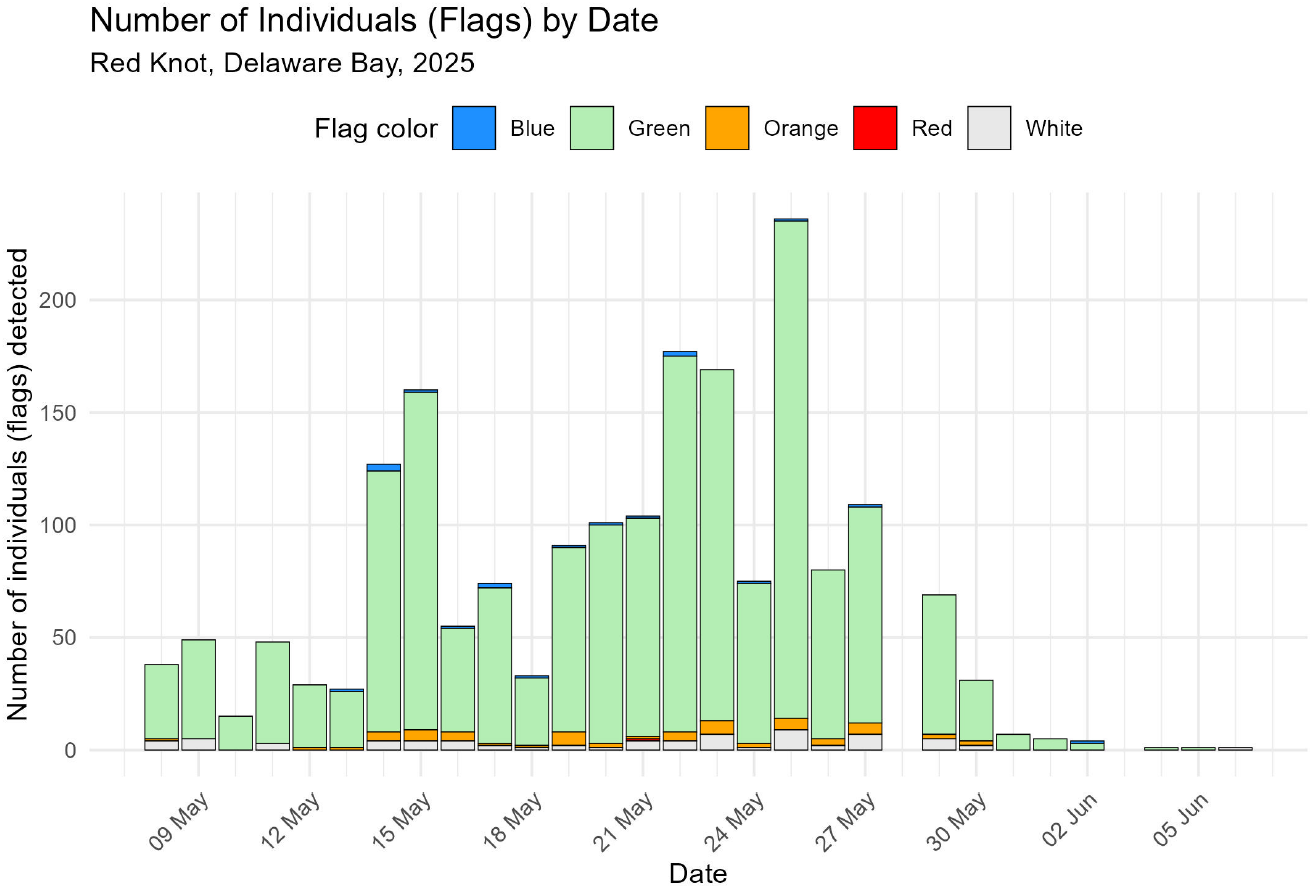
Number of unique leg-flags (individuals) detected each day in 2025. Colors correspond to countries in the Western Hemisphere according to the Pan American Shorebird Program (Howes, Béraud, and Drolet-Gratton, 2016): blue = Brazil and Paraguay; green = USA; orange = Argentina and Uruguay; red = Chile; white = Canada.

**Figure 2.**
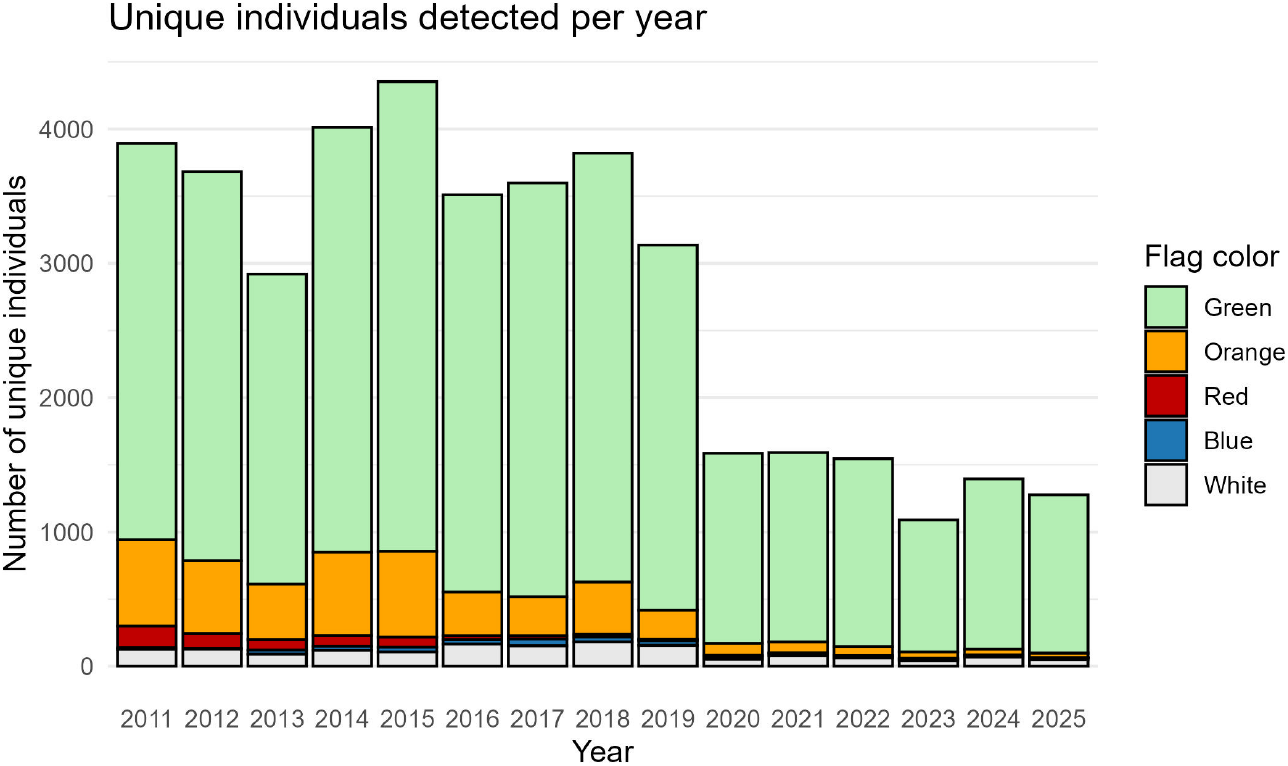
Number of unique leg-flags (individual Red Knots) detected at Delaware Bay each year since 2011. Colors correspond to countries in the Western Hemisphere according to the Pan American Shorebird Program (Howes, Béraud, and Drolet-Gratton, 2016): blue = Brazil and Paraguay; green = USA; orange = Argentina and Uruguay; red = Chile; white = Canada. Data are available from http://www.bandedbirds.org/.

Migrants typically depart Delaware Bay in late May (Niles et al., 2008) and in 5 out of 15 years since 2011, there was little data collected during period 10 (4-6 June; J. Lyons, personal observation, 2025-09-01). In these five years, period 10 was not included in the analysis due to sparse data. In 2025 there were enough resighting observations for analysis in all 10 sampling periods, which span 7 May to 6 June (Fig. 1; Table 1). Although the amount of data from the last two periods in 2025 (1-6 June) was relatively small, the observations were consistent with the sampling plan and suitable for analysis (Appendix B). Field observers also collected 590 marked ratio scan samples in 2025, 327 in Delaware and 263 in New Jersey (Appendix D). The flagged proportion declined to 4.5% in 2025 (95% CI: 3.4%–5.5%; Appendix D), below 2024 values and consistent with a continuing decline in relative abundance of flagged birds compared to unflagged.

### 3.2 Migration ecology and superpopulation estimate for 2025

The majority of arrivals occurred early in the season; by 21 May 2025, 84% of the stopover population was present, and only limited arrivals followed (Fig. 3a). Arrivals were concentrated between 15 and 21 May 2025, although approximately 18% of the stopover population arrived before 12 May, i.e. before period 2. In 2025, departure probability was relatively low (∼ 12%) before 21 May (Fig. 3b). From 18-30 May, departure probability steadily increased, peaking near the end of the month, indicating steady turnover in the population after 24 May. Probability of resighting in 2025 ranged from approximately 14% to 25% between 9 and 27 May, relatively low compared to most years since 2011 and has declined since 2011 (J. Lyons, U.S. Geological Survey, unpublished data, 2025-09-01). The resighting probability of dark green flags in 2025 was not significantly different than other flags (slope: −0.077, 95% credible −0.266 to 0.683; Fig. 3c).

**Figure 3.**
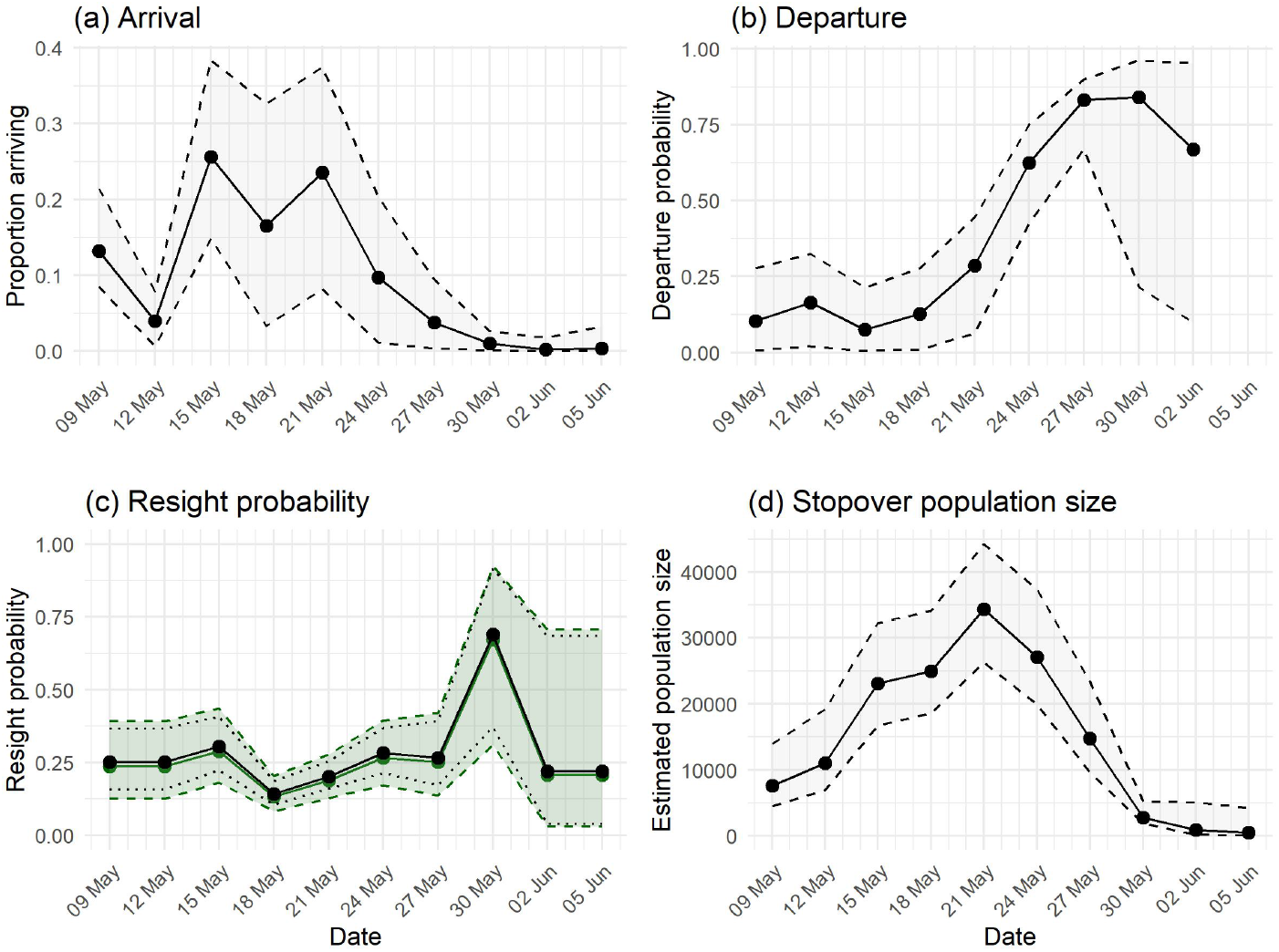
Jolly-Seber model parameter estimates from a mark-resight study of Red Knot (*C. c. rufa*) at Delaware Bay in 2025: (a) proportion of stopover population arriving at Delaware Bay, (b) stopover departure probability, (c) probability of resighting, and (d) time-specific population size estimates. Dates on the x-axis indicate the mid-point of three-day sampling occasions (Table 1). Dashed lines and shading indicate a 95% Bayesian credible interval. The model included two parameters for probability of resighting (c), one for dark green flags and one for other flag colors combined.

The passage population size estimate for 2025 was 54,044 (95% CI: 47,926 – 61,928; Table 2). The time-specific stopover population estimate at the first sampling occasion (9 May) was 7,634 (95% CI 4,525 – 13,991) and increased steadily to a peak of 34,376 by 21 May (Fig. 3d). The stopover population then steadily declined until 30 May and remained below ∼3,000 thereafter.

**Table 2:** Red Knot (*C. c. rufa*) stopover (passage) population estimated using mark-resight methods during northward migration at Delaware Bay. The mark-resight estimate of stopover (passage) population, *N*^∗^, accounts for population turnover during migration. CI = Bayesian credible interval and CV = coefficient of variation.

| Year | Stopover population ($N^*$) | 95% CI $N^*$ | CV |
| --- | --- | --- | --- |
| 2011 | 43,570 | (40,880–46,570) | 0.03 |
| 2012 | 44,100 | (41,860–46,790) | 0.03 |
| 2013 | 48,955 | (39,119–63,130) | 0.12 |
| 2014 | 44,010 | (41,900–46,310) | 0.03 |
| 2015 | 60,727 | (55,568–68,732) | 0.06 |
| 2016 | 47,254 | (44,873–50,574) | 0.03 |
| 2017 | 49,405<sup>[a]</sup> | (46,368–53,109) | 0.03 |
| 2018 | 45,221 | (42,568–49,508) | 0.04 |
| 2019 | 45,133 | (42,269–48,393) | 0.03 |
| 2020 | 40,444 | (33,627–49,966) | 0.10 |
| 2021 | 42,271 | (35,948–55,210) | 0.11 |
| 2022 | 39,800 | (35,013–51,355) | 0.10 |
| 2023 | 39,361 | (33,724–47,556) | 0.09 |
| 2024 | 46,127 | (39,286–57,799) | 0.11 |
| 2025 | 54,044 | (47,926–61,928) | 0.07 |

<sup>a</sup> Data from observers with greater than average misread rate were not included in the analysis in 2017.

Applying the Lok et al. (2019) method, the mean stopover duration in 2025 was 8.6 days (95% CI: 7.9–9.5). Although numerically similar to 2023–2024 estimates (9.0 – 9.2 days; Lyons, 2024), the 2025 value reflects a slight increase when accounting for methodological differences. Nevertheless, estimates from 2023–2025 suggest shorter stays than those from 2019–2021, when stopover duration ranged from 10.3 to 12.1 days (Lyons, 2023).

### 3.3 Timing of arrivals since 2011

The date at which cumulative arrivals reached 50% of the stopover population varied little across the 15 years of the study, ranging from occasion 3 (2023) to occasion 5 (2011 and 2021; Fig. 4). The mean 50% arrival date across all years was 4.2 occasions (range: 3.4-5.0), i.e., approximately 18 May (range: 16–21 May). Linear regression revealed no significant temporal trend in 50% arrival dates over the study period (slope = −0.031 occasions per year, R^2^ = 0.106, p = 0.235), indicating that the timing by which half the population arrived each year remained relatively stable. The earliest timing of arrivals was found in 2019 and 2023. In contrast, 2011 and 2021 showed the latest 50% arrival dates (median = occasion 5, [21 May]). The proportion of years considered “early” based on the stated definition (50% threshold before 18 May [occasion 4]) was 6.7% (1 of 15 years: 2023).

**Figure 4.**
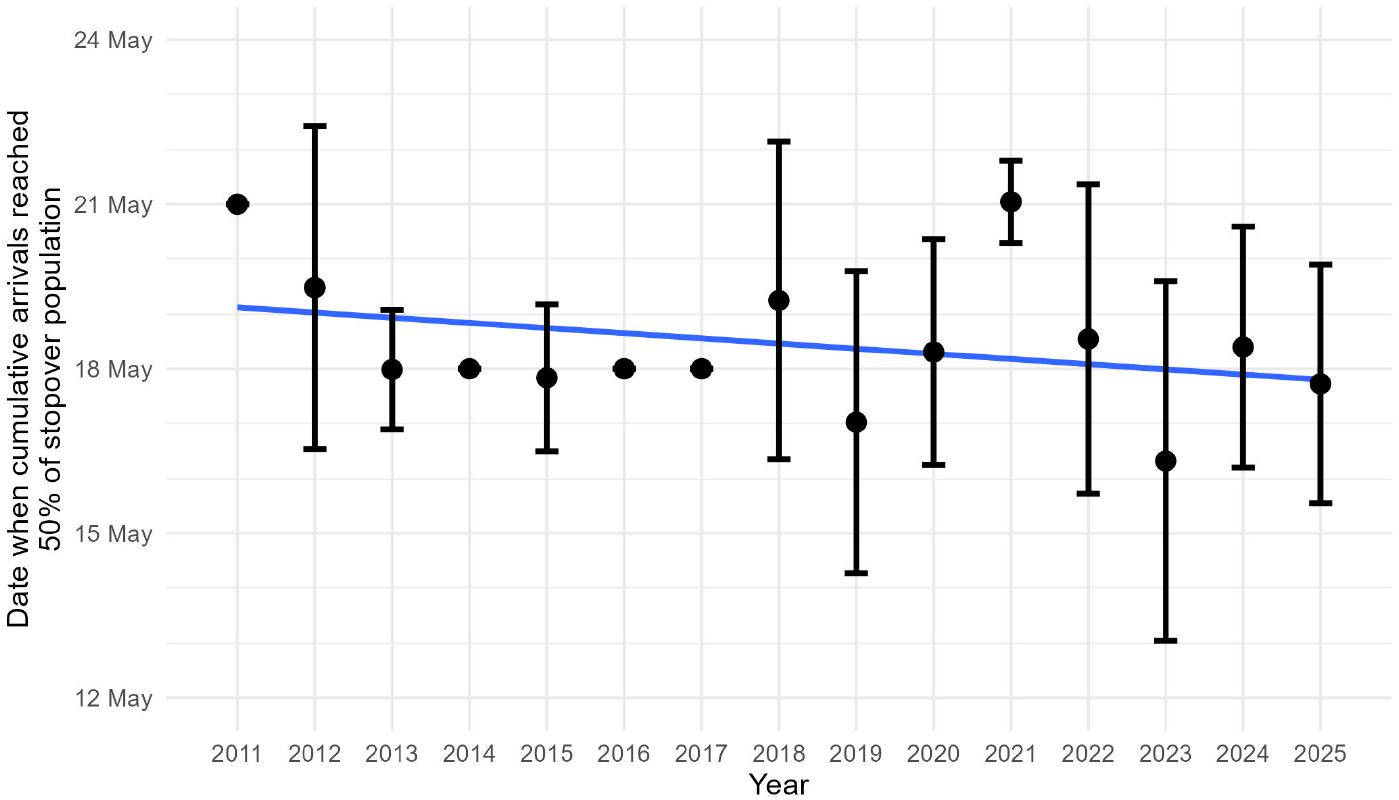
Arrival timing of Red Knots (*C. c. rufa*) at Delaware Bay each year between 2011 and 2025. Dates on the y-axis correspond to sampling occasions 2–6. Points show the occasion when cumulative arrivals reached 50% of the stopover population. Arrival timing is derived from arrival (entry) probability of the Jolly-Seber model. Error bars indicate a 95% Bayesian credible interval for the date when cumulative arrivals reached 50%. Solid line shows the fit of a linear model (slope [change per year] = −0.03, p = 0.24).

## 4 Discussion

By jointly estimating arrival, departure, detection, and total passage population size, the JS model provides robust measures of stopover dynamics at Delaware Bay, which are essential for informing adaptive management of horseshoe crabs and supporting conservation of the Red Knot stopover population.

In 2025, the passage population was estimated at 54,044 Red Knots, 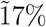 higher than 2024 and only the second estimate since 2011 to exceed 50,000; however, overlapping credible intervals with recent years (generally ∼34,000–58,000 during 2020–2024) indicate no clear directional trend, but rather interannual variability within the long-term range. Producing this annual estimate meets a core goal of the study—providing the ARM framework with up-to-date population inputs to evaluate whether current horseshoe crab harvest strategies maintain adequate foraging opportunities for knots at Delaware Bay.

Overall, the timing of arrivals has not changed significantly since 2011; 50% of the stopover population arrives by 18 May on average (Fig. 4). The analysis revealed little inter-annual variability in arrival timing, with the difference between earliest (2023) and latest (2011, 2021) 50% arrival dates spanning two occasions, representing approximately 6 days within the migration season. The results of the trend analysis should be interpreted with caution because the linear model does not account for the variation in the annual estimates of 50% arrival date; the posterior medians (n = 15) were treated as observed data in the regression analysis. Nevertheless, regression diagnostics did not indicate lack of fit, suggesting that this model presents a reasonable exploratory analysis.

The flagged proportion in 2025 was the lowest percentage since data collection began in 2011 and continues a declining trend in the percentage of the population with leg flags. Historically, this percentage has been close to 10% and was as high as 9.6% (95% CI: 8.8%–10.3%) as recently as 2020 (Lyons, 2020). The fraction of the stopover population with leg flags has been declining at approximately 0.02% per year since 2011 (J. Lyons, U.S. Geological Survey, unpublished data, 2025-09-01). The number of orange flags in the stopover population has declined from approximately 25% in 2011 to 3% in 2025 (Fig. 2); reductions in banding effort in Argentina and Uruguay may explain in part the declining proportion of flagged Red Knots.

The stopover duration of Red Knots at Delaware Bay was slightly higher in 2025 than recent years (2020-2024), but remains lower than the early years of the study (10-12 days in 2013-2016). This suggests that knots may be spending less time in the study area than previous years. Our estimates of stopover duration have limited power to discern small changes in stopover time, however, because they are based on the number of time steps in the mark-resight model that individuals are estimated to remain in the study area. They are coarse measures, in this study, because we use three-day periods as the time step, rather than daily data, reducing the precision of stopover estimates.

By simultaneously estimating arrival and departure schedules, resight probability, and passage abundance, the Jolly–Seber framework delivers integrated, decision-relevant measures of Delaware Bay stopover dynamics. These outputs provide a rigorous basis for adjusting crab harvest within the ARM framework. Mark-resight-derived stopover metrics translate monitoring data into policy-relevant evidence needed to sustain the Delaware Bay stopover and the *rufa* Red Knot population.

## Acknowledgments

This study relied on the dedication and expertise of volunteers and staff from multiple agencies and non-governmental organizations. We thank the scores of volunteers in Delaware and New Jersey who collected mark-resight data in 2025. We are grateful to Katherine Christie (Delaware Division of Fish and Wildlife), Kathy Clark and Bill Pitts (New Jersey Department of Environmental Protection, Division of Fish and Wildlife, Endangered & Nongame Species Program), Stephanie Feigin (Wildlife Restoration Partnerships), Jeannine Parvin, Lena Usyk (bandedbirds.org), and numerous volunteers in Delaware and New Jersey for data entry and data management. We thank A. Royle for guidance on Bayesian statistical modeling approaches and code development. Two peer reviewers provided helpful comments on the manuscript.

## Appendix

### A Study Area

**Table A1:**
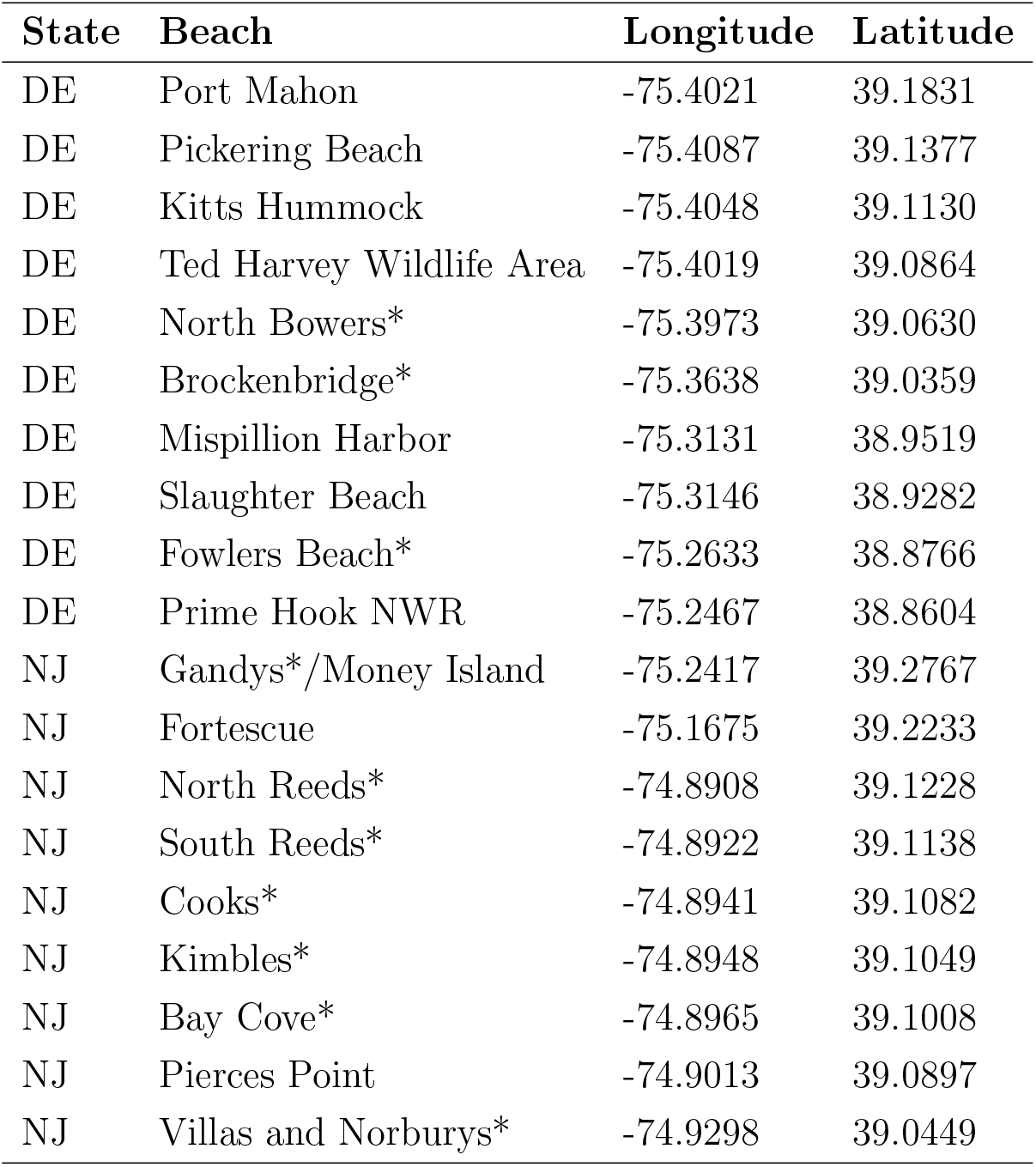
Locations around Delaware Bay, USA, where mark-resight surveys were conducted to estimate Red Knot (*C. c. rufa*) stopover population size during northward migration in 2025. DE = Delaware; NJ = New Jersey; NWR = National Wildlife Refuge. Beach names reflect locally used survey locations. Official geographic names were verified against the U.S. Geological Survey Geographic Names Information System (GNIS) where available. *Locations without GNIS-recognized beach names are identified using locally used names and precise latitude/longitude coordinates.

### B Mark-resight Data Summary

**Table B1:** Summary (m-array) of Red Knot (*C. c. rufa*) mark-resight data from Delaware Bay, USA, 2025. Releases is the number of flags detected at each sample. NR = never resighted.

| Sample | Dates | Releases | Next resighted at sample |  |  |  |  |  |  |  |  | NR |
| --- | --- | --- | --- | --- | --- | --- | --- | --- | --- | --- | --- | --- |
|  |  |  | 2 | 3 | 4 | 5 | 6 | 7 | 8 | 9 | 10 |  |
| 1 | 8-10 May | 84 | 19 | 13 | 3 | 5 | 8 | 0 | 0 | 0 | 0 | 36 |
| 2 | 11-13 May | 100 |  | 26 | 4 | 12 | 4 | 2 | 0 | 0 | 0 | 52 |
| 3 | 14-16 May | 303 |  |  | 43 | 38 | 34 | 8 | 5 | 0 | 0 | 175 |
| 4 | 17-19 May | 188 |  |  |  | 31 | 30 | 3 | 1 | 0 | 0 | 123 |
| 5 | 20-22 May | 355 |  |  |  |  | 64 | 20 | 10 | 0 | 0 | 261 |
| 6 | 23-25 May | 428 |  |  |  |  |  | 45 | 10 | 0 | 0 | 374 |
| 7 | 26-28 May | 177 |  |  |  |  |  |  | 20 | 0 | 0 | 157 |
| 8 | 29-31 May | 96 |  |  |  |  |  |  |  | 5 | 0 | 91 |
| 9 | 1-3 June | 6 |  |  |  |  |  |  |  |  | 1 | 5 |

### C Statistical Methods to Estimate Stopover Population Size Using Mark-Resight Data and Counts of Marked Birds

We converted the observations of marked Red Knots into encounter histories, one for each bird, and analyzed the encounter histories with a Jolly-Seber (JS) model (Jolly, 1965; Seber, 1965). The JS model includes parameters for recruitment (*β*), survival (*ϕ*), and capture (*p*) probabilities; in the context of a mark-resight study at a migration stopover site, these parameters are interpreted as probability of arrival to the study area, stopover persistence, and resighting, respectively. Stopover persistence is defined as the probability that a bird present at time *t* remains at the study area until time *t* + 1.

The Crosbie and Manly (1985) and Schwarz and Arnason (1996) formulation of the JS model also includes a parameter for superpopulation size, which in our approach to mark-resight inferences for stopover populations is an estimate of the marked (leg-flagged) population size.

We chose to use three-day periods, rather than days, as the sampling interval for the JS model given logistical constraints on complete sampling of the study area; multiple observations of the same individual in a given three-day period were combined for analysis. A summary (m-array) of the mark-resight data is presented in Appendix B.

We made inference from a fully time-dependent model; arrival, persistence, and resight probabilities were allowed to vary with sampling period [*β*_*t*_, *ϕ*_*t*_, *p*_*t*_]. In this model, we set *p*_1_ = *p*_2_ and *p*_*K*−1_ = *p*_*K*_ (where *K* is the number of samples) because not all parameters are estimable in the fully time-dependent model (Jolly, 1965).

We followed the methods of Royle and Dorazio (2008, Chapter 10) and Kéry and Schaub (2012, Chapter 10) to fit the JS model using the restricted occupancy formulation. Royle and Dorazio (2008) use a state-space formulation of the JS model with parameter-expanded data augmentation. For parameter-expanded data augmentation, we augmented the observed encounter histories with all-zero encounter histories (*n* = 2, 000) representing potential recruits that were not detected (Royle and Dorazio, 2012).

According to the Pan American Shorebird Program (Howes, Béraud, and Drolet-Gratton, 2016), researchers in the U.S. deploy green engraved leg flags. In 2014, researchers switched from using light green flags to dark green flags, which many observers found more difficult to read in the field, as shown by Tucker, McGowan, Nuse, et al. (2023). Therefore we modeled two detection probabilities, one for dark green flags and one for all other flag colors:

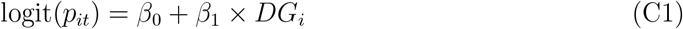

where *β*_0_ is the logit of the average resighting probability of all non-dark-green flags, *β*_1_ is the effect of dark green flag color on resighting probability, and DG = 1 for dark green flags and 0 otherwise.

We followed Lyons et al. (2016) to combine the JS model with a binomial model for the counts of marked and unmarked birds in an integrated Bayesian analysis. Briefly, the counts of marked birds (*m*_*s*_) in the scan samples are modeled as a binomial random variable:

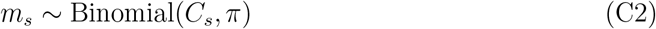

where *m*_*s*_ is the number of marked birds in scan sample *s, C*_*s*_ is the number of birds checked for marks in scan sample *s*, and *π* is the proportion of the population that is marked. Total stopover population size 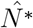 is estimated using

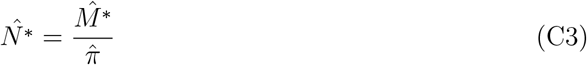

where 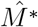 is the estimate of marked birds from the JS model and 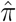 is the proportion of the population that is marked (from Eq. C2). Estimates of marked subpopulation sizes at each resighting occasion *t*, 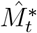, are available as derived parameters in the analysis. An estimate of population size at each mark-resight sampling occasion, 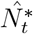, is calculated using 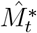 and 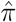 as in equation C3.

To better account for the random nature of the arrival of marked birds and addition of new marks during the season, we used a time-specific model for proportion with marks in place of equation C2 above:

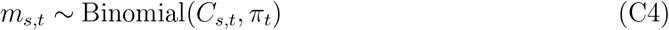

for *s* = 1, …, *n*_samples_ and *t* = 1, …, *n*_occasions_

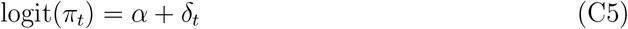

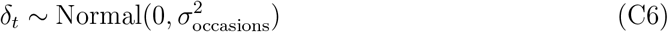

where *π*_*t*_ is the time-specific proportion of the population that is marked, and *δ*_*t*_ is a random effect for time of sample *s*. Total stopover population size 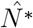 was estimated by summing time-specific arrivals of marked birds to the stopover (*B*_*t*_) and expanding to include unmarked birds using estimates of proportion marked:

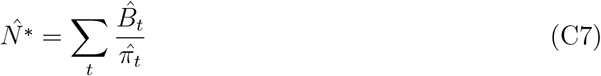

Time-specific arrivals of marked birds are estimated from the JS model using 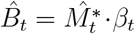, where *β*_*t*_ is the estimated fraction of the population arriving at time *t*.

### D Marked-ratio Scan Samples

**Figure D1.**
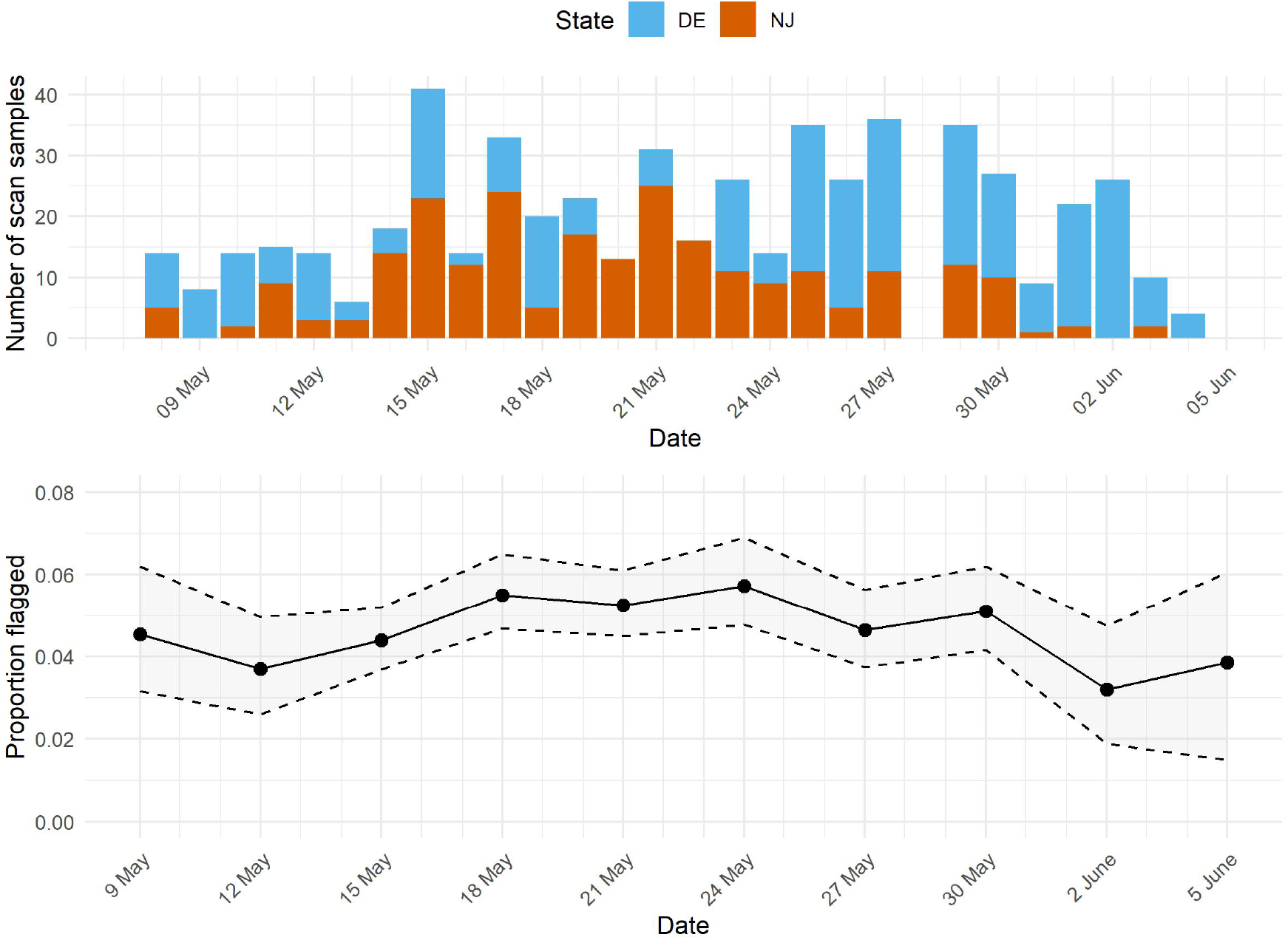
(top) Number of marked-ratio scan samples (n = 590) collected at Delaware Bay in 2025 by field crews in Delaware (blue, n = 327 scan samples) and New Jersey (orange, n = 263 scan samples). (bottom) Estimated proportion of the Red Knot (*C. c. rufa*) stopover population at Delaware Bay carrying leg flags in 2025 (overall average and 95% Bayesian credible interval: 0.045 [0.034, 0.055]). The marked proportion was estimated from marked-ratio scan samples for each three-day sampling period (Table 1) and the generalized linear mixed model described in Appendix C. Solid and dashed lines are estimated median proportion marked and 95% Bayesian credible interval, respectively.

